# Kinome profiling allows examination and prediction of kinase inhibitor cardiotoxicity

**DOI:** 10.64898/2026.04.03.716310

**Authors:** Jimmy S. Tabet, Chinmaya U. Joisa, Brian C. Jensen, Shawn M. Gomez

## Abstract

**Background:** Despite improved cancer outcomes with kinase inhibitors (KIs), their cardiotoxicity remains a significant clinical challenge. Current approaches to predict and prevent KI-induced cardiac adverse events (CAEs) are limited by an incomplete understanding of underlying mechanisms, including the contribution of off-target kinase engagement.

**Objectives:** To establish links between kinase inhibition profiles and cardiotoxic phenotypes using empirical proteomic data, and to leverage these profiles in machine learning (ML) models capable of predicting KI cardiotoxicity.

**Methods:** We curated a database connecting kinome-wide target binding profiles of FDA-approved KIs (n=44) with documented incidence rates of six distinct CAEs. Binding profiles were derived from unbiased chemoproteomics and used to assess associations between KI selectivity, specific kinase targets, and CAEs. Profiles were further used to develop ML models to predict CAE risk, with SHAP-based model interpretation applied to identify cardiotoxicity-associated kinases.

**Results:** KI promiscuity was not a significant predictor of cardiotoxicity across all six CAEs. Frequency analysis revealed that kinases including RET, PDGFRB, and DDR1 are recurrently inhibited across CAE-linked compounds, with nearly all identified as off-targets not annotated by the FDA. Network and pathway enrichment analyses supported a systems-level model in which cardiotoxicity arises from coordinated disruption of cardiac-relevant signaling networks. ML models achieved 66–84% cross-validated accuracy (ROC-AUC 0.75–0.8) across CAE endpoints, with SHAP analysis identifying PDGFRB, EGFR, and MEK1/2 among the most predictive kinases.

**Conclusions:** Proteomic kinome profiling combined with machine learning provides a mechanistically grounded framework for predicting KI cardiotoxicity and supports off-target-aware drug design to minimize cardiovascular risk.

## Introduction

While cytotoxic chemotherapy remains a mainstay in cancer treatment, targeted therapies have led to improved outcomes in a variety of cancers(1, 2). Examples include imatinib (Gleevec) for chronic myelogenous leukemia, crizotinib and other anaplastic lymphoma kinase (ALK) inhibitors for non-small-cell lung cancers, and trastuzumab and lapatinib for ERBB2/HER2 amplified breast cancers (3–8). These successes, coupled with relatively favorable adverse effect profiles have fueled the rapid expansion of targeted therapy development over the last two decades (9, 10).

Since the introduction of imatinib, kinases have emerged as a dominant focus for targeted oncology drug development (11–14). Comprising a 500-member enzyme family, kinases regulate a broad range of cellular processes including growth, proliferation, differentiation, motility, and apoptosis, through phosphorylation of downstream substrates (15). As evidence of the kinome’s integral role in this wide array of functions, dysregulation of one or more kinases is directly implicated in numerous pathologies, most notably numerous types of cancer (16–18). This central role in disease, combined with the feasibility to modulate kinase activity through small molecule targeted inhibition, has positioned kinase inhibitors (KIs) as one of the fastest growing drug classes, with 100 inhibitors having now been approved by the FDA (19– 21) and over 1300 trials actively enrolling patients (clinicaltrials.gov).

Despite initial optimism that targeted therapies would carry more favorable cardiovascular profiles than cytotoxic agents such as anthracyclines, KI-based therapies have not fully realized this promise. Clinical manifestations of KI-induced cardiovascular damage span a spectrum from reversible cardiomyopathy to hypertension, QT interval prolongation, arrhythmia, myocardial infarction, overt heart failure, and sudden cardiac death (22, 23). Though the type and severity of toxicity varies across agent and molecular target, dozens of KIs have recognized adverse cardiovascular effects (summarized in (24, 25)), and in CML alone, KIs were implicated as the probable driver in 83% of cardiovascular events (26). The true burden of these effects is consistently underestimated in clinical trials due to enrollment of low-risk patients and limited efforts at detection – often only becoming evident in post-marketing or pharmacovigilance studies (27, 28). As cancer survivorship improves and the patient population ages, the clinical significance of KI cardiotoxicity will only grow (29, 30).

The consequences extend beyond patient outcomes: cardiac toxicity is a leading cause of drug attrition, estimated to account for 14–45% of clinical trial withdrawals across multiple eras of drug development (31–33). There is thus a pressing need for tools capable of predicting cardiotoxicity earlier, ideally in the preclinical space, both to guide investment toward cardiosafe compounds and to reduce downstream cardiovascular morbidity.

A key challenge is that the mechanisms underlying KI cardiotoxicity remain poorly defined. KIs vary widely in selectivity, and both on-target inhibition of cardioprotective kinases and off-target polypharmacology contribute to observed toxicity. Because most KIs target the highly conserved ATP-binding pocket, off-target kinase engagement is common and often uncharacterized. Furthermore, the intuitive assumption that broader, more “promiscuous” inhibitors carry greater cardiac risk has not been consistently borne out, suggesting that cardiotoxicity is driven more by which kinases are engaged than by how many.

Prior modeling efforts to predict KI cardiotoxicity have drawn on chemical structure, transcriptomic profiles, and large drug safety databases, and have demonstrated promising results (34–38). However, a critical gap remains: no existing approach has directly incorporated empirical, proteome-wide data on how KIs bind and perturb the kinome. Advances in chemical proteomics and more particularly, multiplex inhibitor bead/mass spectrometry (MIB/MS) and Kinobeads approaches, now make it possible to profile the binding of hundreds of kinases to a given inhibitor in a single experiment, providing an unbiased, quantitative map of the full target landscape (39, 40). This data type is uniquely positioned to capture the off-target interactions that drive cardiotoxicity but are invisible to target-annotation-based approaches.

Here, we integrate a comprehensive kinome-wide binding database for FDA-approved KIs with clinically documented cardiac adverse event (CAE) incidence rates to address three aims: (1) evaluate whether target promiscuity correlates with CAE incidence; (2) identify specific kinase targets disproportionately associated with cardiotoxic outcomes; and (3) develop and interpret machine learning models that predict CAE risk from global kinome binding profiles. Together, these analyses establish a novel, proteomics-driven framework for preclinical cardiotoxicity prediction and rational design of safer kinase-targeted therapies.

## Methods

### Curation of a cardiac adverse event (CAE) incidence rate database for FDA approved kinase inhibitors in cancer

Data defining clinical cardiotoxicity incidence of FDA-approved small molecule kinase inhibitors was taken from Jin et al.(23), where the cardiac adverse event (CAE) incidence rates for each approved inhibitor as of 2020 were annotated for six common CAEs: QT interval prolongation, heart failure, ischemia or myocardial infarction, arrhythmia, left-ventricular dysfunction, and hypertension (23). The incidence rates of these CAEs were annotated as three groups: 1) no reported event, 2) reported event but with an incidence rate of less than 10%, and 3) an incidence rate exceeding 10%. The final 38 cardiotoxic drugs from Jin et al. were supplemented with 17 more drugs we noted as “negative controls” as they were SMKIs approved as of 2020 per Roskoski (41), but were not mentioned in Jin et al.’s review. These 17 drugs received a label of “no reported event” for all CAEs. Note that one of the cardiotoxic drugs from Jin et al., Abemaciclib, was not associated with any of the six common CAEs but was noted to have “cardiovascular malformations and variations.” For this study exploring the six common CAEs, Abemaciclib was included as a negative control. In total, we accumulated 55 drugs with information across six CAEs (see Table 2).

### Connecting the quantitative kinase inhibitor target-ome to CAE incidence rates

As described previously (42, 43) a database of 229 drugs and their kinome-wide binding profiles was curated from the study of Klaeger et al.,(39) containing for each drug the kinase target space, selectivity, and full dose-response characteristics, spanning nearly across the entire kinome. This dataset was then connected to the CAE incidence rate dataset via drug synonym matching per PubChem. Eleven drugs (20%) were not matched/not profiled by Klaeger et al, resulting in a final dataset of 44 drugs with associated binding/activity profiles, shown in bold in Table 2. Of these 44 drugs, 12 (27%) were negative controls with no reported cardiotoxicity (including Abemaciclib). In order to establish the definition of a drug target using the Klaeger et al. derived inhibition percentage, a cutoff of >80% inhibited was used.

### CAE incidence rate modeling

Machine learning models using the kinome profile-CAE incidence rate were built using a random 5-fold cross validation strategy with hyperparameter tuning. We first binarized the incidence rate variable as follows: “reported event” and “>10% incidence” were combined into the positive class, indicating cardiotoxicity, while “no reported event” remained as the negative class. We then aimed to predict if a given inhibitor would have any incidence of each of the six cardiotoxic events. By training six separate models, we were able to develop CAE-specific inferences. We compared the performance of multiple model types surveying three major categories of machine learning algorithms: linear models, tree-based models, and neural networks. Linear models included logistic regression (LR), tree-based models included random forest (RF) and gradient boosting using XGBoost38 (XGB) and LightGBM39 (LGBM), while neural networks included simple feed forward multi-layer perceptron models (MLP) and a transformer-based foundation model (TabPFN40)(44–48). Hyperparameter tuning (Table 1) was performed over 20 iterations of randomized search to extract optimal performance from each model type. Tuning and model training were implemented using the scikit-learn package in Python(48). The best performing model was chosen based on mean accuracy across the five folds, with ties favoring the model with the smallest standard deviation. Model interpretation was performed using SHAP42 (SHapley Additive exPlanations) in Python. SHAP is a model-agnostic interpretation method derived from cooperative game theory that quantifies the contribution of each feature to a model’s prediction (49).

**Table 1.**
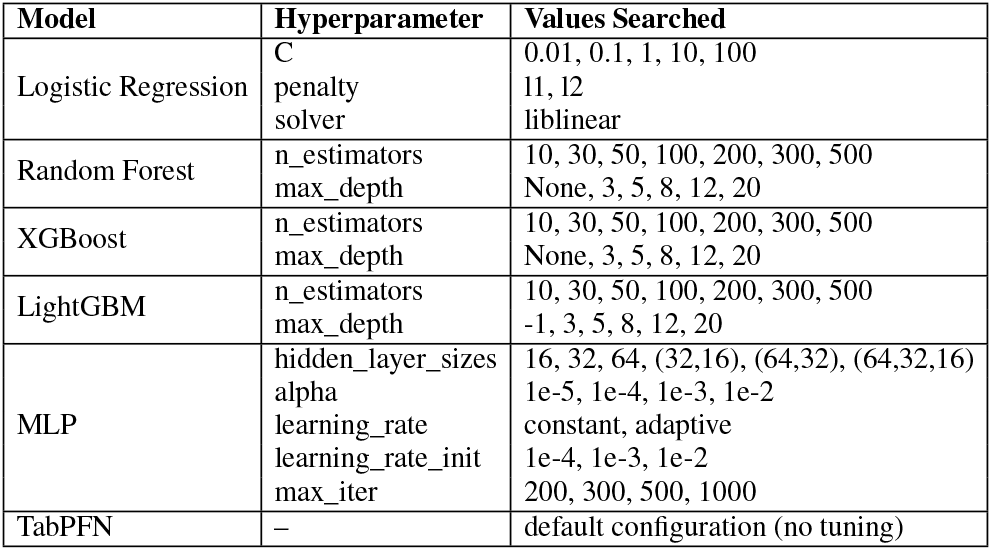
Hyperparameter search space for different models. Hyperparameters were tuned using 20 iterations of randomized search over the ranges listed in the table. The best configuration was selected based on 5-fold cross validation performance.

## Results

### Defining kinase inhibitor proteomic kinome binding profiles

We first sought to connect both our CAE-linked and control (no strong link to CAEs) inhibitors defined in Table 2 with their corresponding proteomic binding profiles. For each FDA-approved compound, we utilized the proteomic profiles characterized in Klaeger et al. (39) (see Methods), and plot inhibitors and their targets as a heatmap, a subset of which are shown in Figure 1 (full heatmap available as Supplementary File S1). The heatmap values indicate experimentally measured percent inhibition/binding of a kinase for each KI (concentration 1 µM). Providing a global view of inhibitor targets, this heatmap shows both the on-and off-target binding spectrum for each inhibitor as well as the broad variability of targeting across inhibitors. Importantly, there is no clear property that distinguishes cardiotoxic vs control inhibitors, suggesting that there are less-obvious underlying drug targeting patterns that drive CAEs. We further note that the FDA labeling typically assigns a limited number of kinase targets for each drug, while the proteomic data reveals broader inhibition across the kinome, uncovering dozens of additional kinase targets. This discrepancy underscores a gap between regulatory (FDA) and functional (proteomic) target annotation, revealing potentially targets that should be assessed for their role in CAEs.

**Table 2.**
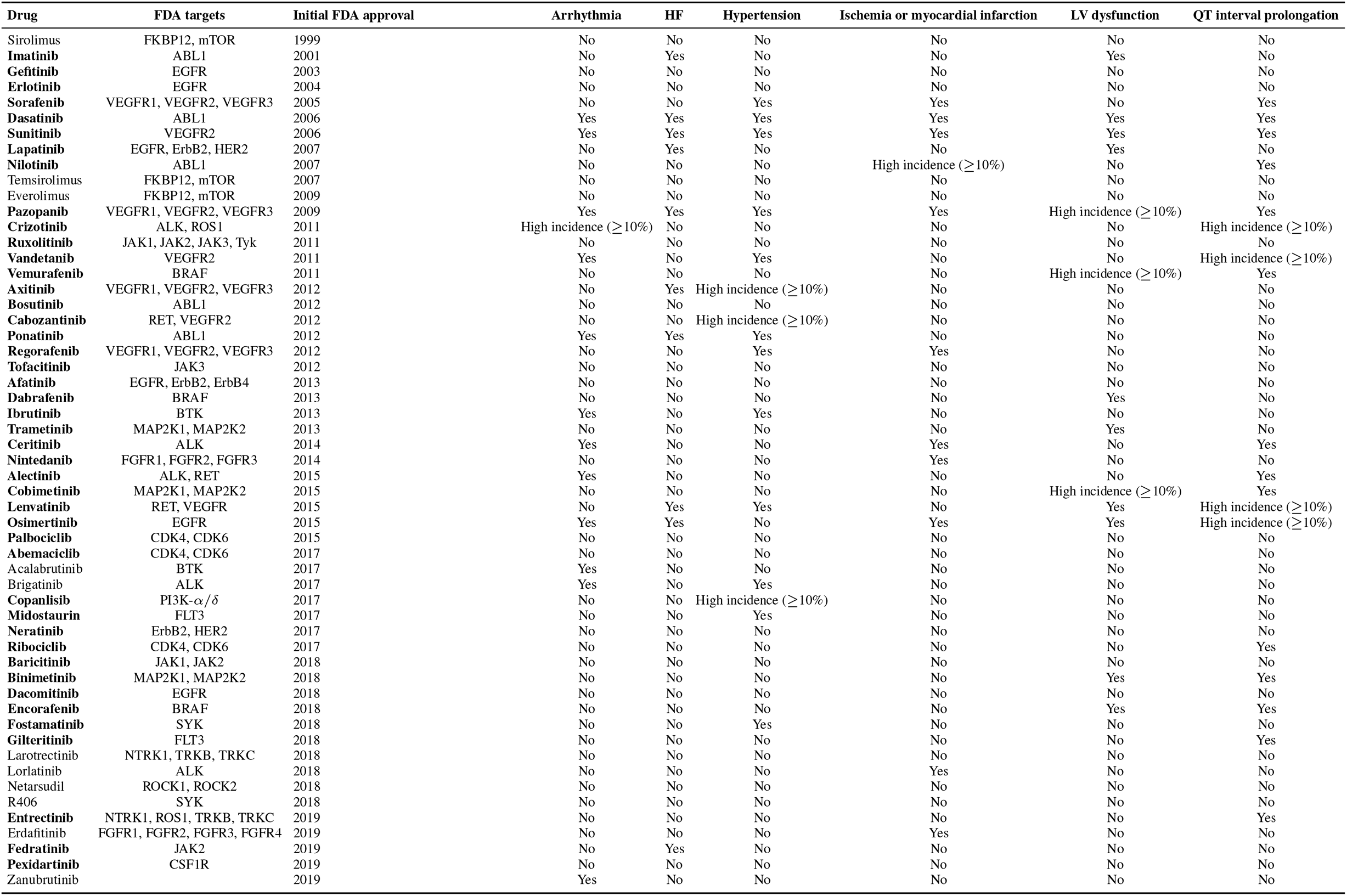
Cardiotoxicity of small molecule kinase inhibitors (SMKIs) in clinical use. Derived from the comprehensive review by Luo et al, this table supplements their findings of FDA-approved cardiotoxic kinase inhibitors (KIs) with 17 negative controls (i.e. KIs that did not have reported cardiotoxicities). In total, 55 drugs were compiled, though 11 could not be matched with proteomic-derived kinome inhibition profiles. The remaining 44 drugs (shown in bold in Table 1) were used to train our predictive machine learning models.

**Fig. 1.**
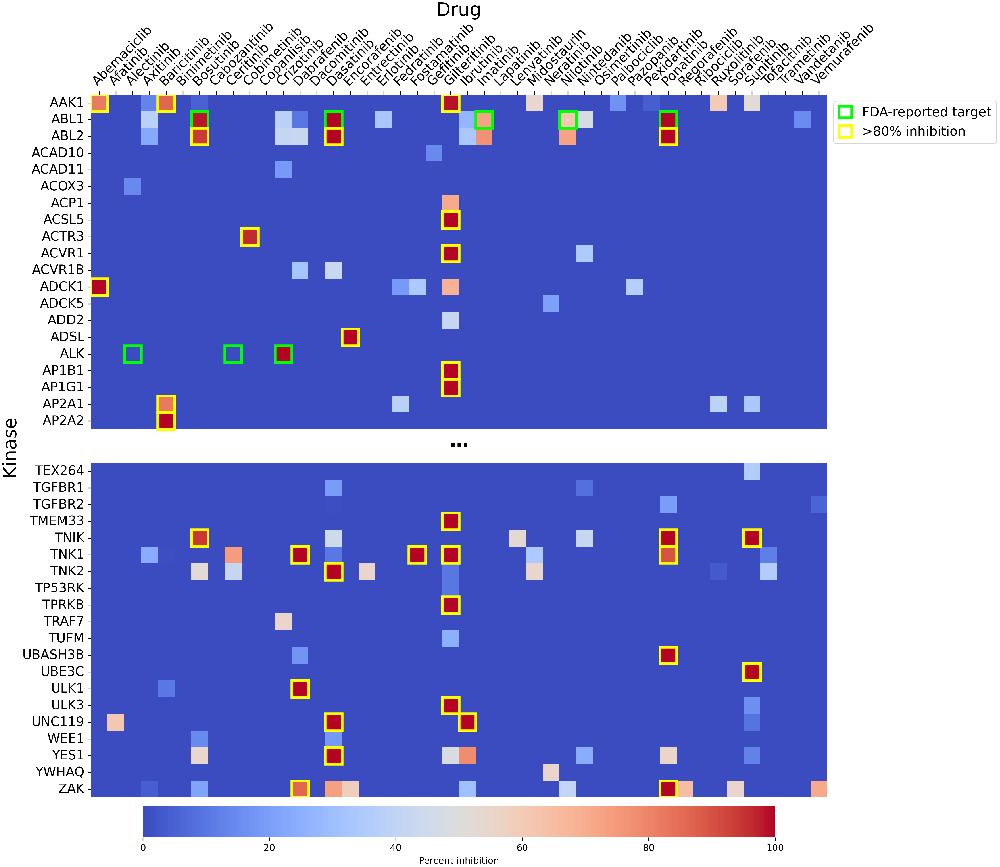
Heatmap of kinome inhibition profiles reveals discrepancies between FDA-reported kinase inhibitor targets and proteomic-derived targets. Heatmap values indicate experimentally measured percent inhibition of kinases for each KI (concentration 1 µM). Green boxes highlight FDA-reported primary targets of the KIs, while the yellow boxes indicate proteomic-derived targets as defined by greater than 80% reduction in relative intensity. Full, high-resolution kinome inhibition state heatmap available as Supplementary File S1.

### KI target promiscuity does not fully explain incidence rates of CAEs

Having connected the complete binding profile of KIs to their CAE incidence rates, we can interrogate the intuitive hypothesis that a “dirty” or promiscuous inhibitor will have more cardiac adverse effects associated with it than does a more selective inhibitor. To investigate this possibility, we created a promiscuity score for all KIs in our dataset by counting the number of kinases inhibited by at least 80% by each inhibitor. This threshold was chosen to focus on inhibitors having the greatest impact on kinase behavior.

For each of the six CAE categories, we compared promiscuity scores between the “No Reported Event,” “Reported Event,” and “>10% Incidence” rates (Figure 2). Across all CAEs, no strongly statistically significant difference in mean promiscuity was observed between the three incidences (Kruskal-Wallis p > 0.04 for all six CAEs). Of note, LV dysfunction represented an exception with weak statistical significance (p=0.041) indicating a difference in mean promiscuity, with a measured increase in promiscuity associated with the Reported Event case. In addition, we investigated the sensitivity of these results to changing the degree of inhibition from 80% to 20% and found that there were again no statistical differences across categories as a function of inhibitor promiscuity, including for LV dysfunction (data not shown). Overall, these findings suggest that broad inhibition of the kinome alone does not explain the incidence of CAEs. This result also supports a model in which cardiotoxicity is not driven by nonspecific kinase inhibition burden, but rather by selective inhibition of particular kinases or kinase subnetworks critical to cardiac function.

**Fig. 2.**
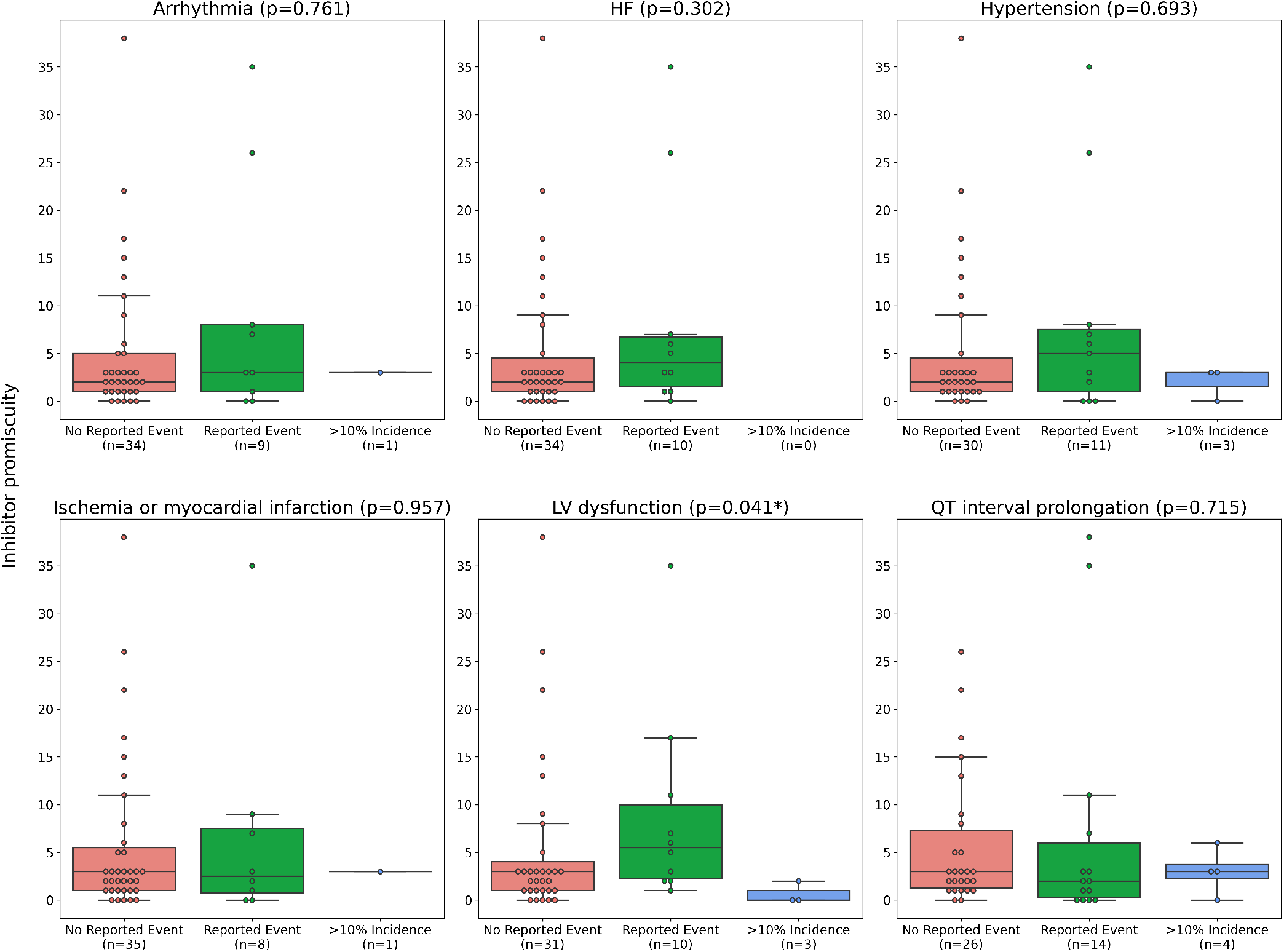
Inhibitor promiscuity does not explain cardiotoxicity. Promiscuity of an inhibitor, defined as the number of kinase targets inhibited by at least 80% by the compound, is shown for each CAE. The median KI promiscuity score was 3, though values ranged to >35 for more promiscuous inhibitors. Incidences were reported as either No Reported Event, Reported Event, or >10% Incidence.

### Comprehensive analysis of all off-and on-targets reveals frequently occurring targets associated with cardiotoxic events

We then examined both off-and on-target interactions of identified inhibitors to identify specific kinases that are frequently inhibited across drugs linked with CAEs. As shown in Figure 3, this analysis reveals both conserved and CAE-specific patterns of kinase targeting. Several kinases (RET, PDGFRB, DDR1) are recurrently inhibited across all CAEs, suggesting potential central roles for them, or the signaling networks within which they function, in cardiotoxic effects. These kinases tend to have larger node sizes in Figure 3, signifying that many drugs inhibit them and supporting their importance in shared toxicity signals. Conversely, other kinases demonstrate more exclusive associations with certain CAEs, such as LV dysfunction and QT interval prolongation. Importantly, we note that the majority of kinases identified in Figure 3 as frequently targeted by CAE-inducing inhibitors are not the primary or designed target for these drugs. Instead, nearly all kinases shown in Figure 3 are off-target hits as identified through proteomic profiling. Collectively, these results suggest that cardiotoxicity reflects overlapping yet distinct inhibition patterns, perhaps heavily influenced by off-target effects, and supporting the need for a more network-oriented view of inhibitor-associated cardiotoxicity to reveal potential mechanistic relationships among targeted kinases.

**Fig. 3.**
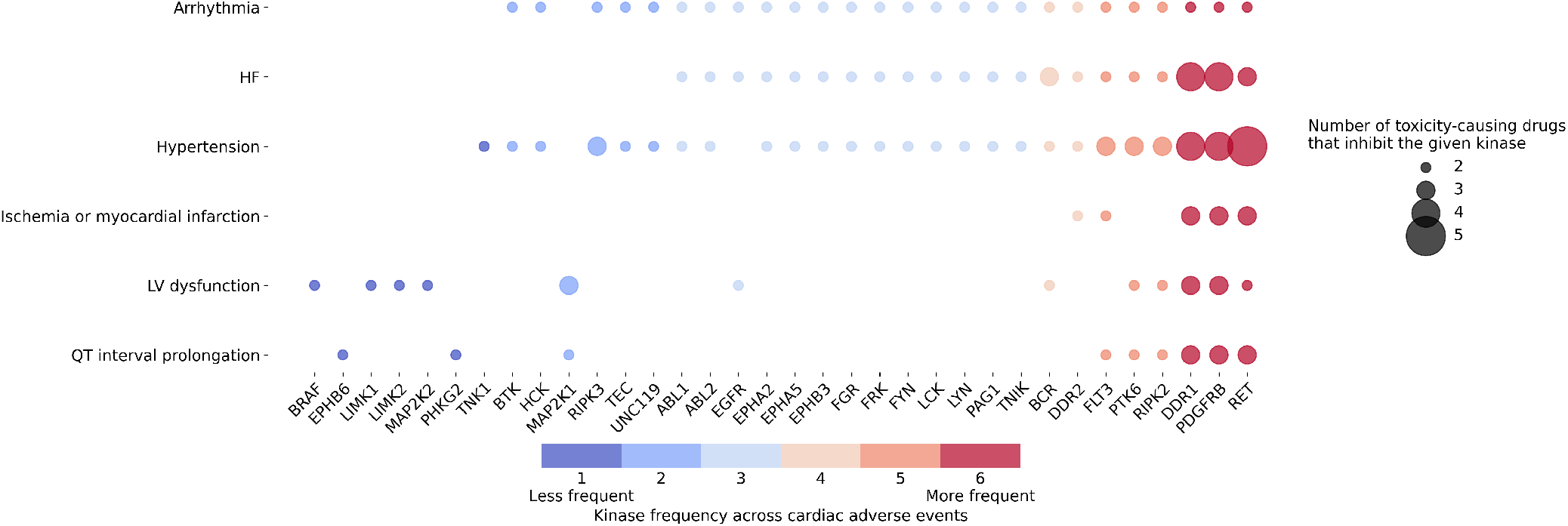
Kinase inhibition frequency varies across cardiac adverse events highlighting shared kinase targets and CAE-specific inhibition patterns. Shown are kinase targets (x-axis) that are inhibited by two or more kinase inhibitors associated with a specific CAE (y-axis). Node size represents the number of toxicity-causing drugs inhibiting the given kinase. Kinases are arranged from left to right according to their frequency across CAEs, with those recurrently inhibited across all six CAEs shown in red on the right.

### Network interpretation

Building upon the kinase inhibition frequency analysis (Figure 3), frequently inhibited kinases associated with representative CAEs were mapped into protein-protein interaction networks using the HIPPIE protein interaction database and KEGG signaling pathways, with representative networks shown for hypertension and LV dysfunction shown in Figure 4 (50, 51). Nodes for the protein interaction networks represent the CAE-specific kinases identified in the frequency analysis, with edges denoting experimentally supported protein interactions. As in Figure 3, node size in the networks reflects the number of toxicity-causing drugs inhibiting a given kinase. These networks clearly demonstrate the often-extensive degree of interaction occurring between CAE-implicated kinases.

**Fig. 4.**
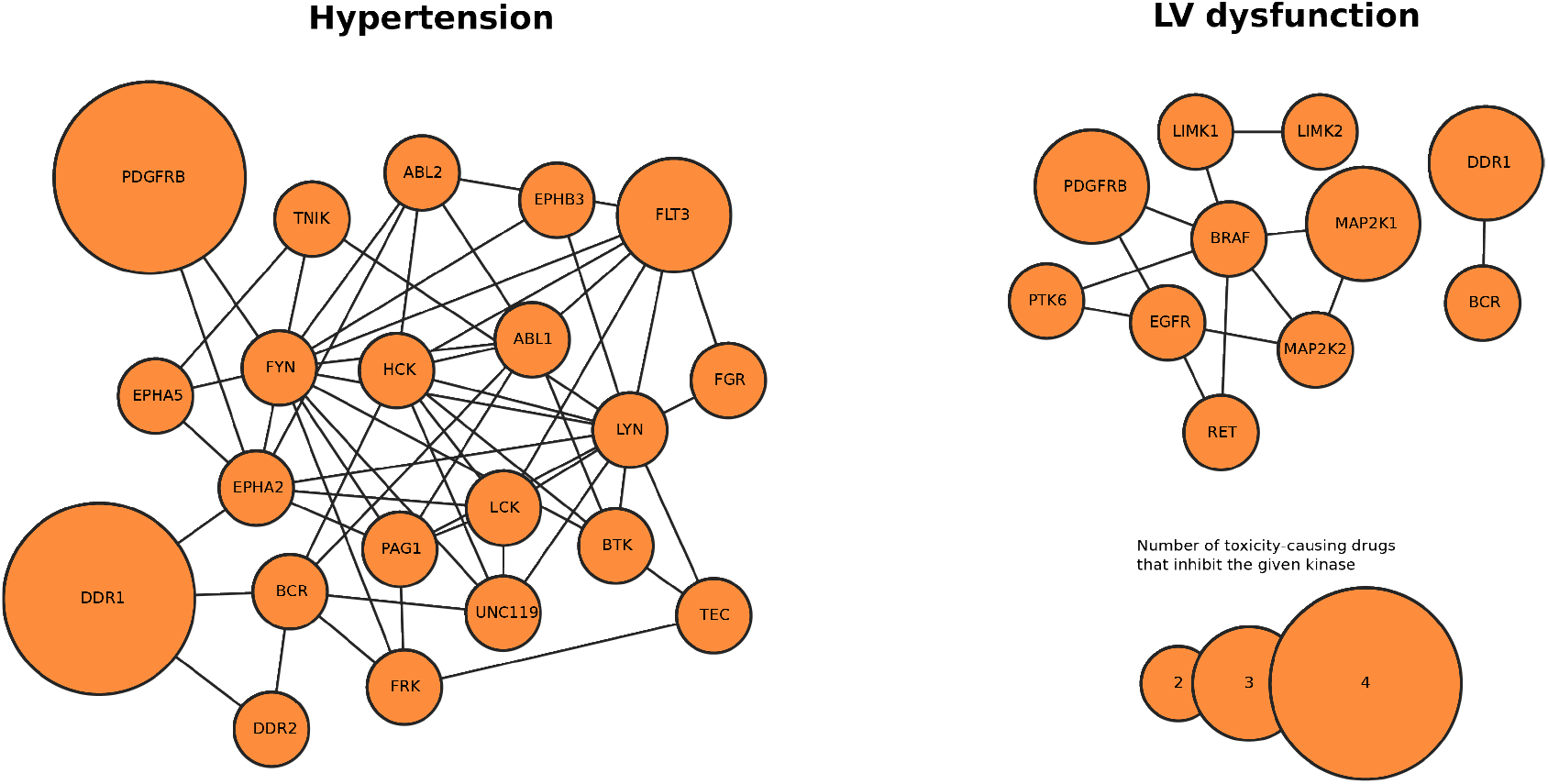
Frequently inhibited kinases across CAEs form interaction networks enriched for cardiac-relevant KEGG pathways. Example protein interaction networks for hypertension (left, most enriched KEGG pathway: axon guidance) and LV dysfunction (right, most enriched KEGG pathway: regulation of actin cytoskeleton) are shown. Nodes represent the CAE-specific kinases identified in the frequency analysis and edges denote experimentally supported protein interactions from HIPPIE. Node size reflects the number of toxicity-causing drugs inhibiting a given kinase. See Supplementary Figure S2 and S3 for hypertension and LV dysfunction KEGG pathways, respectively. (a)5-fold cross-validation performance of multiple machine learning algorithms trained to predict the occurrence of CAEs from kinome inhibition profiles. (b)SHAP feature importance analysis for the best performing model for each CAE

To further characterize the biological context of these interactions, over-representation analysis (ORA) was performed using the g:Profiler functional profiling service to identify enriched biological pathways among the frequently inhibited kinases(52). ORA evaluates whether genes from the input list occur within annotated biological pathways more frequently than expected by chance. Statistical significance is assessed using cumulative hypergeometric probability with multiple testing correction applied using g:Profiler’s custom g:SCS algorithm. For each CAE, kinase gene lists derived from the frequency analysis were tested for enrichment against the KEGG pathway database. In the KEGG pathway diagrams (Supplementary Figure S2 and S3), kinases identified from frequency analysis are outlined in red indicating where frequently inhibited targets fall within established signaling cascades. Notably, these enriched pathways correspond to biologically relevant processes including axon guidance in the case of hypertension and the actin cytoskeleton for LV dysfunction, consistent with known contributors to cardiac dysfunction. These results extend the frequency-based findings and support the systems-level model indicating that cardiotoxicity arises from coordinated disruption of interconnected signaling networks embedded within cardiac-relevant pathways.

### Global kinome profiles of KIs are predictive of CAE incidence through machine learning

After examining individual cardiotoxic targets, we examined whether proteomic kinome profiling data could be used to predict CAE incidence. We tested the predictive ability of multiple machine learning algorithms including linear (logistic regression), tree-based (random forest, XGBoost, LightGBM)(44, 45, 47), and neural network-based (MLP, TabPFN)(46, 48) approaches, to connect the KI profiles their downstream CAE incidence rates (see Methods; Figure 5a). Models underwent 5-fold cross validation (CV), with individual model architectures undergoing hyperparameter tuning to extract maximum performance. Looking across models and CAEs, cross validation accuracy demonstrated very strong performance, with model accuracy ranging from 66% for QT interval prolongation to 84% for hypertension. However, no single architecture clearly stood out as broadly superior as a predictive platform across CAEs. Excluding QT interval prolongation, the best model for a given CAE performed at or above 80% in CV accuracy, with ROC AUC scores typically in the range of 0.75-0.8.

**Fig. 5.**
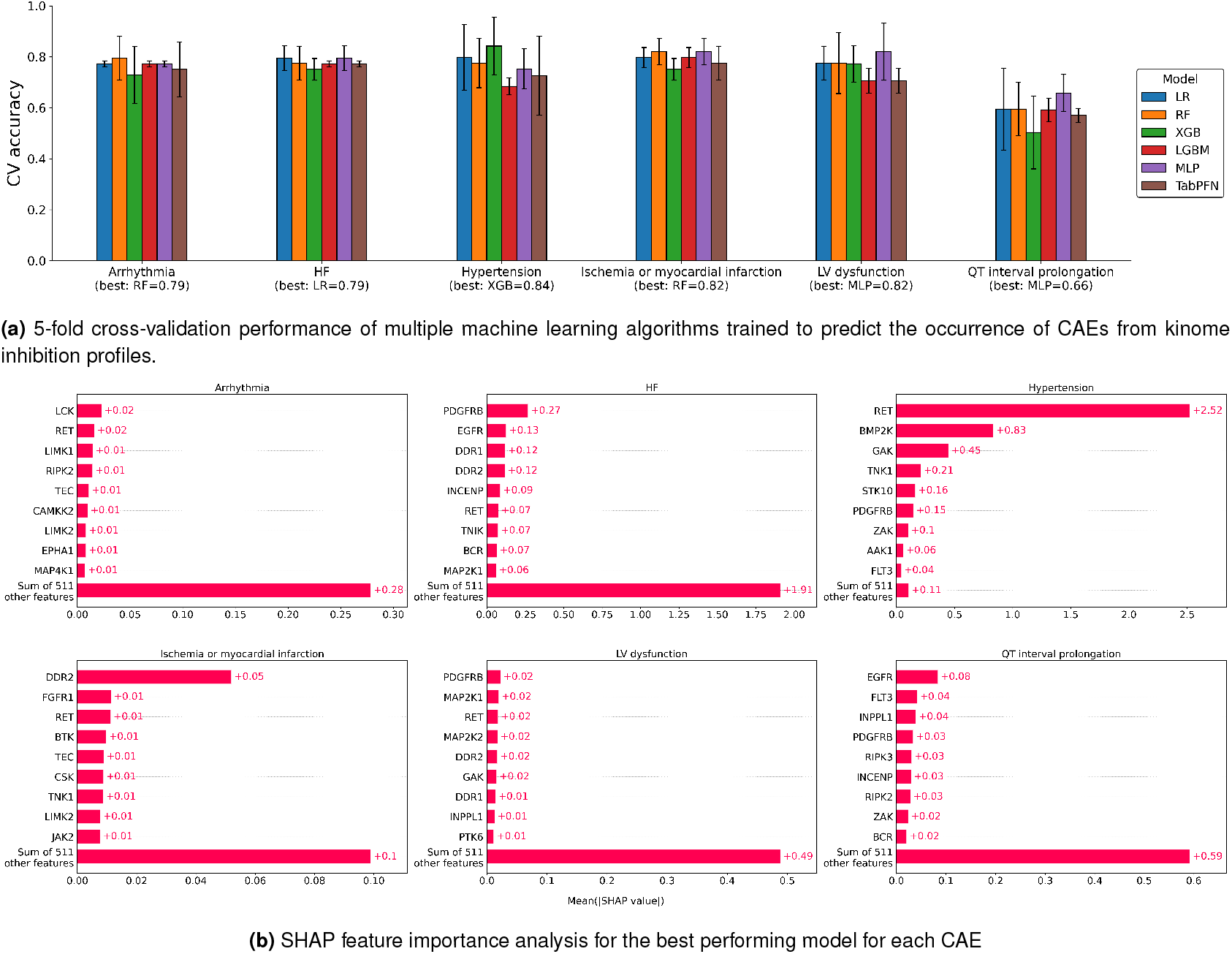
Kinome inhibition profiles alone enable prediction of multiple cardiac adverse events.

As the models used for prediction are interpretable, we then sought to identify those kinase targets that were most predictive of a CAE risk within our models through SHAP analysis (Figure 5b). SHAP (SHapley Additive exPlanations) is a model-agnositc interpretation method derived from cooperative game theory that quantifies the contribution of each feature to a model’s prediction(49). In this context, SHAP values estimate each kinase’s contribution to predicting whether a drug is associated with a given CAE. Higher mean absolute SHAP values indicate features (i.e. kinases) that are most influential to the models’ predictions, allowing identification of the most predictive inhibition signals.

Across endpoints, the models prioritized distinct but biologically coherent kinase sets, supporting the hypothesis that specific inhibition signatures underlie different cardiotoxic phenotypes. Many of the highly-ranked kinases are already recognized for their involvement, directly or indirectly, with CAEs, including PDGFRB, EGFR and MEK1/2. These kinases are often implicated in myocardial stress signaling, hypertrophic remodeling, vascular smooth muscle proliferation, inflammatory signaling, and cardiomyocyte survival. In addition, this analysis also identifies kinases that are less, or perhaps only marginally linked with cardiac dysfunction, such as RET, LCK, LIMK1, TEC, TNK1, DDR1, and FLT3. More commonly studied in the context of immune signaling, cytoskeletal regulation, or oncogenic pathways, these kinases emerge as influential predictors in different CAE endpoints. Together, these results suggest that current KIs may induce cardiotoxicity directly through both impacting canonical cardiac signaling pathways as well as through previously under-appreciated kinase interactions.

## Discussion

To our knowledge, this study presents the first systematic integration of empirical, proteome-wide kinase binding data with clinical cardiac adverse event incidence rates across the FDA-approved KI landscape. By leveraging unbiased proteomic profiling data quantifying binding across hundreds of kinases simultaneously, this work establishes a foundation for understanding and predicting KI-induced cardiotoxicity that goes well beyond what is possible using FDA-annotated primary targets alone. Three principal findings emerge from this work. First, target promiscuity is a poor predictor of CAE risk. Second, cardiotoxicity is disproportionately associated with inhibition of a specific limited set of kinases, the majority of which are off-targets invisible to current regulatory annotation. Third, machine learning models built on global proteomic kinome binding profiles achieve meaningful pre-dictive accuracy for multiple CAE endpoints and reveal biologically coherent, CAE-specific kinase signatures.

### Promiscuity is not the primary driver of KI-induced cardiotoxicity

A relatively common assumption in the field is that broader, more “promiscuous” kinase inhibitors carry proportionally greater toxicity and specifically, greater cardiotoxic risk than more selective agents. Our data do not support this view. With the exception of a single borderline case (LV dysfunction, any reported event), there is no significant difference in kinase inhibitor promiscuity between cardiotoxic and non-cardiotoxic compounds across all six CAE endpoints examined. This is regardless of whether the inhibition threshold was set at 20% or 80%. This null result is also consistent whether toxicity is defined as any reported event or by events exceeding 10% clinical incidence. Left ventricular dysfunction was the partial exception at the 80% cutoff, with a weak association between higher promiscuity and reported events, but this finding did not survive changes to the inhibition threshold definition.

These findings have direct implications for drug development strategies. Attempts to reduce toxicity are often pursued by designing more selective inhibitors, but our analysis suggests this logic is incomplete. A highly selective inhibitor that happens to engage a cardioprotective kinase will carry significant cardiac risk regardless of its clean overall profile, while a moderately promiscuous inhibitor that avoids cardiotoxic targets may be well tolerated. This suggests that identification of key kinases implicated in cardiosafe and cardiotoxic outcomes should be a central component of therapy development, pointing towards more target-aware design strategies rather than selectivity per se as the appropriate lever for reducing cardiac risk.

### Off-target kinase inhibition underlies cardiotoxic risk

Analysis of kinase inhibition frequency across CAE-associated drugs revealed both shared and CAE-specific patterns of target engagement. Kinases such as RET, PDGFRB, and DDR1 were recurrently inhibited across multiple CAE categories, suggesting central roles in shared cardiotoxic mechanisms—or at minimum, membership in signaling subnetworks that regulate cardiac function more broadly. Importantly, nearly all the kinases identified in this analysis are off-targets: they are not among the primary targets formally annotated by the FDA for these compounds. This finding reinforces a key mechanistic point that has been established in individual drug studies but not previously demonstrated at the scale of the FDA-approved KI landscape—that off-target kinase engagement is a dominant, rather than incidental, contributor to KI cardiotoxicity.

The protein-protein interaction network and KEGG pathway analyses provide biological context for these findings. Frequently co-inhibited kinases tend to be highly interconnected, forming dense subnetworks rather than isolated hits, and these subnetworks map onto cardiac-relevant pathways including axon guidance (a known contributor to vascular tone regulation implicated in hypertension) and actin cytoskeletal organization (a pathway fundamental to cardiomyocyte contractility and structure, relevant to LV dysfunction). These observations support a systems-level model in which KI cardiotoxicity arises from coordinated disruption of interconnected kinase networks embedded in cardiac-relevant biology, rather than from inhibition of any single node. Whether single-node or network-level perturbation is the primary driver remains an important open question that motivates future mechanistic work.

### Kinome binding profiles enable machine learning prediction of cardiac adverse events

A central methodological finding in this work is the demonstration that proteomic kinome binding profiles alone can predict CAE incidence with meaningful accuracy. Cross-validated models achieved ROC-AUC scores in the range of 0.75–0.8 and accuracy of 66–84% across endpoints, with the strongest performance for hypertension and weakest for QT interval prolongation. While these results are based on a small data set, the performance levels achieved under these constraints suggest that kinome binding profiles contain genuine signal with respect to cardiac risk. Furthermore, no single model architecture dominated across all endpoints, suggesting that the signal is real but distributed and that ensemble or multi-model strategies may ultimately yield the best practical predictive performance.

Our approach complements and extends prior predictive efforts. Previous models have leveraged chemical structure, gene expression from cardiomyocyte-like cell lines, or large drug safety databases to predict cardiotoxicity(36–38). Importantly, none of these approaches directly incorporated empirical data on kinase-level target engagement. Chemical structure models capture what a drug looks like, not how it behaves at the proteome level. Transcriptomic approaches capture downstream consequences of kinase perturbation, not the perturbation itself, while other approaches infer targets rather than use direct measurements. While our focus solely on KIs and their proteomically-defined targets limits broader generalization, it provides mechanistically grounded input to prediction that is actionable in drug design as well as interpretable, providing much greater insight into potential mechanistic explanations that drive CAEs.

### Model interpretation reveals CAE-specific kinase signatures

SHAP-based interpretation of the predictive models identified distinct, biologically coherent kinase sets associated with each CAE. Many of the highest-ranked kinases are already recognized in the context of cardiac biology. Specifically, PDGFRB, EGFR, and MEK1/2 have established roles in myocardial stress signaling, hypertrophic remodeling, vascular smooth muscle proliferation, inflammatory cascades, and cardiomyocyte survival pathways. Their emergence as influential predictors provides a degree of internal validation for the modeling approach. Equally important are the kinases that are less well established in cardiac contexts including RET, LCK, LIMK1, TEC, TNK1, DDR1, and FLT3. While requiring further investigation, their high feature importance suggests potentially underappreciated roles in specific cardiotoxic phenotypes that merit experimental follow-up.

### Study limitations

Several limitations should be considered when interpreting our findings. First, our analysis relied on a limited set of FDA-approved kinase inhibitors with documented cardiac adverse event rates, which will not fully represent the diverse chemical and pharmacological landscape of KIs in development. Second, the binary classification of cardiac adverse events (present/absent or <10%/>10% incidence) may oversimplify the complex, dose-dependent spectrum of cardiotoxicity observed clinically.

Our kinome profiling data was derived from a single proteomic data type based on the Kinobeads platform(39). While providing consistent and high-quality readouts compared to other in vitro platforms, it cannot fully recapitulate the complex, in vivo kinetics, tissue distribution, and metabolism of these compounds. Additionally, the cellular context of kinase inhibition likely varies between cardiac tissue and other tissues, potentially affecting the translation of our findings to clinical cardiotoxicity.

The retrospective nature of our analysis using published cardiotoxicity data introduces potential confounding factors, including variable monitoring protocols across clinical trials, underreporting of mild cardiac events, and differences in patient populations that may affect cardiotoxicity susceptibility. Our models also do not account for patient-specific factors such as pre-existing cardiovascular conditions, genetic polymorphisms, or drug-drug interactions that may significantly influence individual cardiotoxicity risk.

Furthermore, while our machine learning approach identified potential cardiotoxic kinase targets, the mechanistic validation of these targets through functional studies remains to be completed. Finally, our analysis focused primarily on direct kinase targeting and did not fully explore potential nonkinase off-targets or secondary mechanisms (such as metabolite toxicity or immune-mediated effects) that may contribute to observed cardiotoxicity profiles.

## Conclusions

This study demonstrates that proteomic kinome profiling provides a powerful and mechanistically grounded basis for predicting kinase inhibitor cardiotoxicity. By linking empirical, quantitative binding data across the full kinase target landscape to clinical cardiac adverse event incidence, we find that cardiotoxic risk is determined not by how promiscuously a KI engages the kinome, but by which specific kinases it engages. This observation has direct implications for how cardiosafety should be considered in KI design and development. The proteomic profiling and modeling approach used here offers several practical advantages for translational deployment. First, these assays are scalable, reproducible, and can be applied to novel compounds early in development, and well before clinical cardiotoxicity data are available. Integration with predictive machine learning frameworks could flag high-risk compounds for more intensive cardiac monitoring or reformulation, while identifying specific cardiotoxic off-target kinases to avoid in future design iterations. As the KI landscape continues to expand and the cancer patient population ages and accumulates cardiovascular comorbidities, tools that integrate cardiosafety prediction into the early drug development workflow will become increasingly essential to delivering both efficacious and tolerable therapies to patients.

## Clinical Perspectives

The clinical burden of KI cardiotoxicity is substantial and growing. With over 100 FDA-approved kinase inhibitors now in clinical use, a rapidly expanding development pipeline, and a cancer patient population that is both aging and surviving longer, the cumulative cardiovascular impact of this drug class will only increase. Cardiotoxicity has been estimated to account for 14–45% of clinical trial withdrawals across eras of drug development, and post-marketing data consistently reveal a higher burden than clinical trials suggest. Patients with pre-existing cardiovascular risk factors, increasingly common in the cancer population, are disproportionately vulnerable. There is thus a pressing need for better preclinical tools to identify cardiotoxic liability before it manifests clinically.

This work potentially addresses this need in two complementary ways. First, as a preclinical screening tool, kinome binding profiles of novel or investigational KIs could be rapidly screened against predictive models to flag compounds with elevated cardiac risk before they advance in development. This approach would be most powerful when combined with in vitro cardiac assays, as kinome binding data could identify which specific targets to interrogate mechanistically. Second, as a rational design guide, the identification of specific cardiotoxic kinase associations provides a quantitative basis for optimizing inhibitor selectivity profiles. Rather than optimizing solely for on-target potency, future computational design workflows could incorporate cardiotoxic target avoidance as an explicit constraint. In the broader context of precision oncology, this approach could ultimately be deployed as one layer in a multi-objective treatment selection framework that weighs predicted antitumor efficacy against predicted cardiac risk for individual patients.

## Supplementary Information

### A. Supplementary Files

**Supplementary File S1:** Full kinome inhibition state heatmap, provided as a separate high-resolution PDF file.

### B. Supplementary Figures

**Fig. S2.**
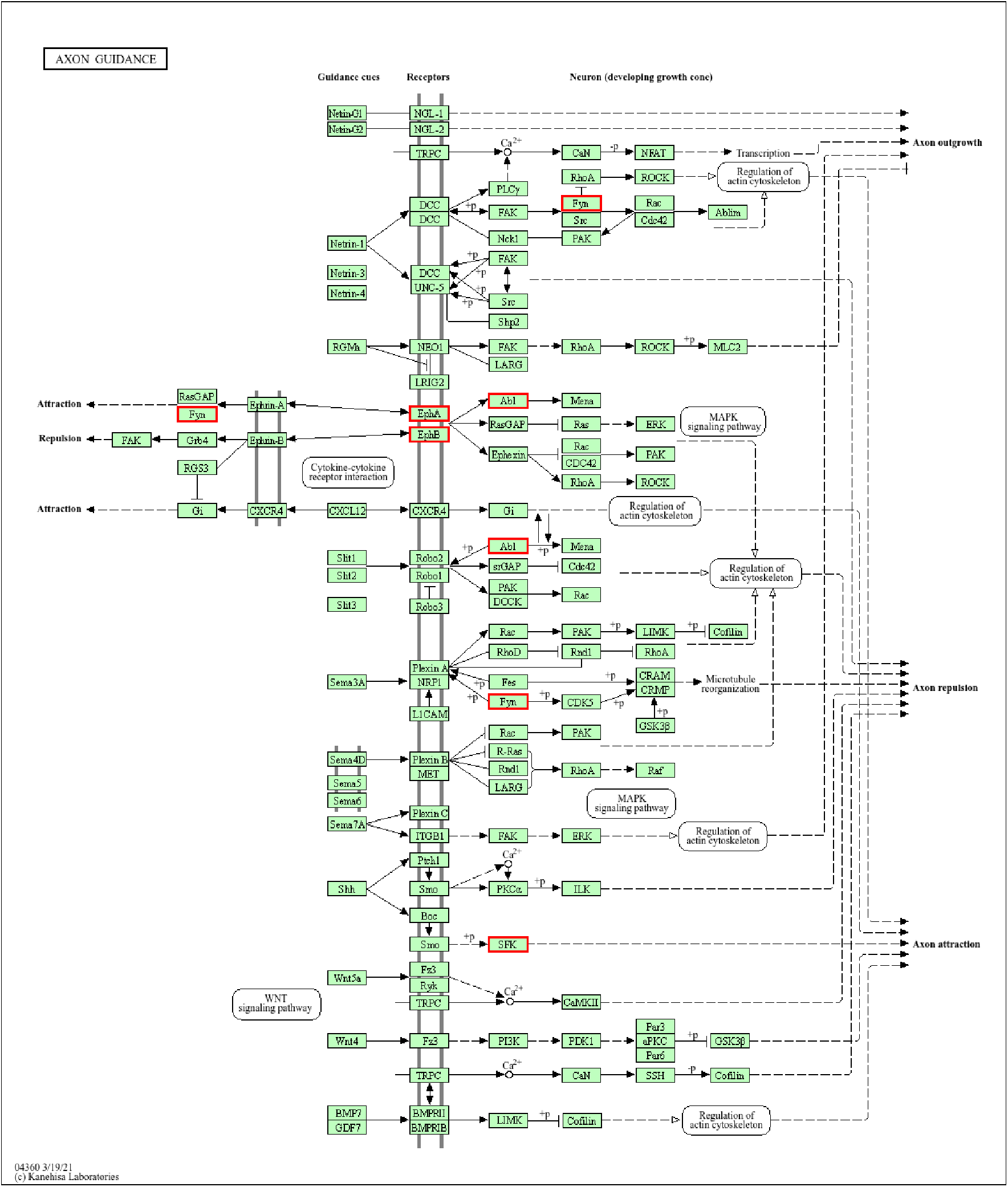
Top KEGG signaling pathway for frequently inhibited hypertension kinases (Figure 4, left): axon guidance (KEGG:04360). P-value: 1.76e-03. Relevant kinases in pathway (highlighted in red): ABL1, EPHA2, EPHA5, EPHB3, FYN.

**Fig. S3.**
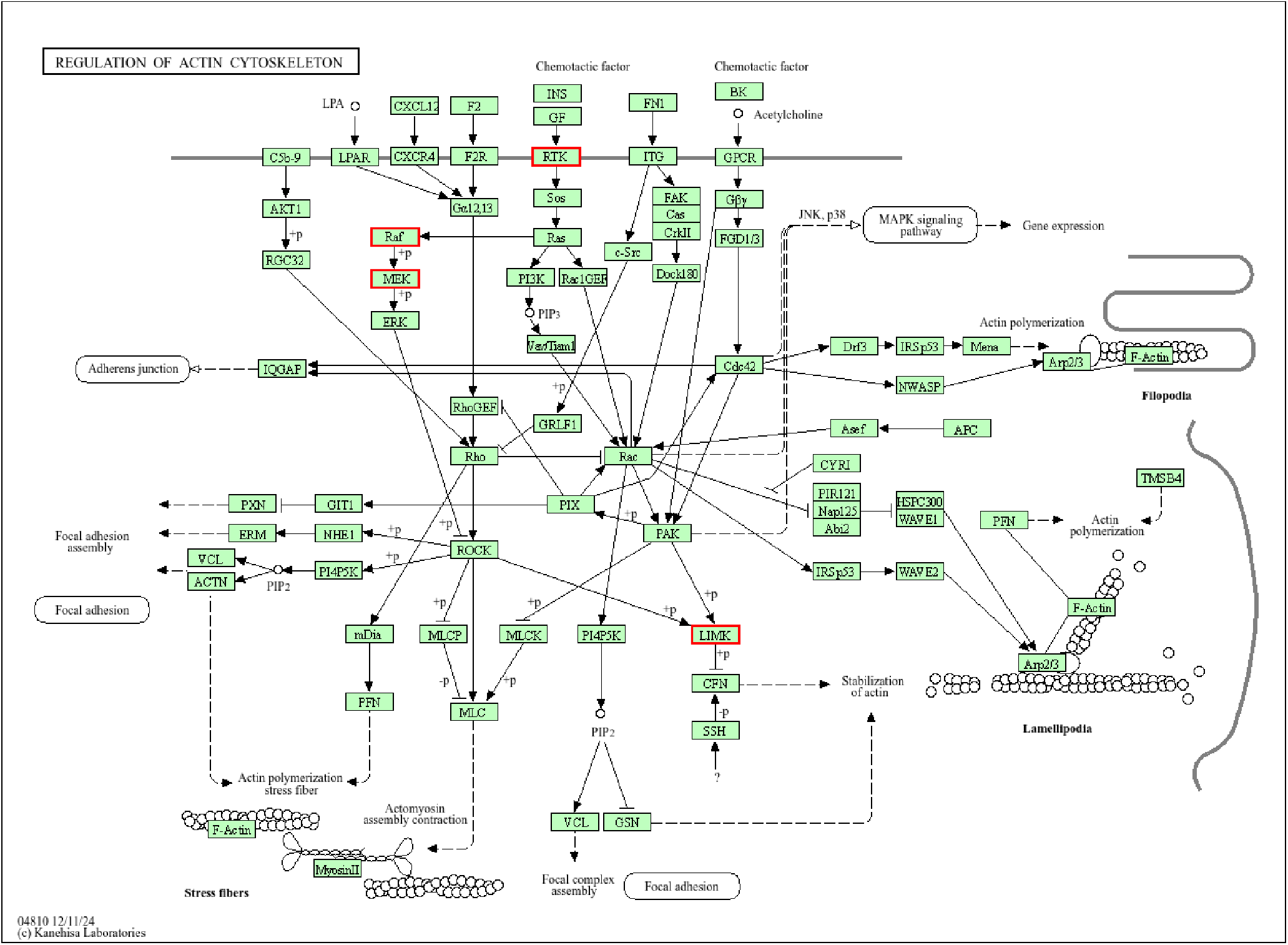
Supplementary Figure S3. Top KEGG signaling pathway for frequently inhibited LV dysfunction kinases (Figure 4, right): regulation of actin cytoskeleton (KEGG:04810). P-value: 5.43e-08. Relevant kinases in pathway (highlighted in red): BRAF, EGFR, LIMK1, LIMK2, MAP2K1, MAP2K2, PDGFRB

