## Supplementary figures and images for "Kinome profiling allows examination and prediction of kinase inhibitor cardiotoxicity"

### Supplementary File S1

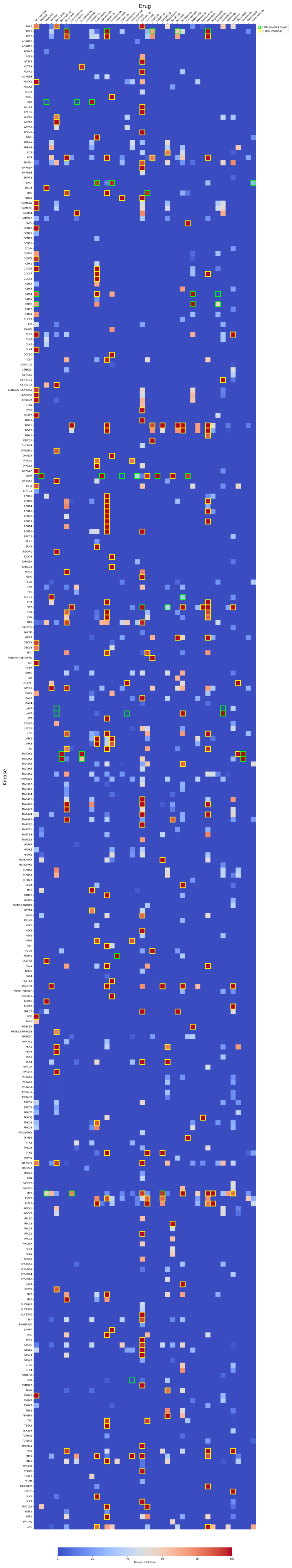
